# Context-dependent hierarchical categorization of human faces: Behavioral and EEG/MEG evidence

**DOI:** 10.1101/2025.02.11.637761

**Authors:** Xuena Wang, Shihui Han

## Abstract

Social categorization of faces provides a key cognitive basis of human behavior and may occur along various dimensions of facial attributes. The present study investigated a potential hierarchical structure of social categorization of faces based on a superordinate (Species) versus a subordinate (Race) level of abstraction of facial attributes. We recorded behavioral performances in a face classification task and found faster responses to the same set of Asian faces when presented alternately with dog faces (a species context) relative to Black faces (a race context). In addition, using a repetition suppression (RS) paradigm, we recorded electroencephalography (EEG) and magnetoencephalography (MEG) signals to Asian faces in the species and race contexts, respectively. Our analyses of the RS effects on EEG/MEG signals to Asian faces revealed that dynamic neural encoding of similarity of Asian faces occurred in the right fusiform gyrus at 140-200 ms and in the left temporoparietal junction at 317-413 ms after stimulus onset in the species context but only in the left temporoparietal junction at 317-413 ms in the race context. These behavioral and EEG/MEG findings unravel the neurocognitive mechanisms of context-dependent social categorization of faces by highlighting its hierarchically organized structure based on different levels of facial attributes.

## Introduction

Humans sort different individuals’ faces into various social categories which enable efficient social interactions and adaptive behaviors. Ample evidence indicates that social categorization of faces may occur spontaneously in terms of age, gender, race, and other attributes (Brewer and Lui, 1989; Kinzler et al., 2010; Todorov et al., 2015; Willis and Todorov, 2006; Zhou et al., 2020; Zhang et al., 2023). Because one face can be sorted into different categories defined by various social attributes, social categorization of faces is challenged by a multiple category problem regarding how perceivers select one out of several available categorical attributes to classify faces (Hugenberg and Sacco, 2008). The present study tested a hypothesis that there is a hierarchy in social categorization of faces such that categorization of faces occurs earlier based on one compared to another level of abstraction of facial attributes. We tested this hypothesis by examining how social contexts influence the dynamic processes of social categorization of faces.

Face perception is thought to start from parallel processing of various facial features that in turn activate corresponding social categories such as race, gender, and age (Freeman and Ambady, 2011; Freeman and Johnson, 2016; Freeman et al., 2020). However, categorization of faces based on different levels of abstraction of facial attributes may be affected by cognitive and social factors that are intrinsically different between faces of different categories. For example, previous event-related potential (ERP) studies found that individuals showed an earlier attentional bias towards Black compared to White faces whereas attention to gender emerged later (Ito and Urland, 2003, 2005), resulting in earlier neural activities that differentiated faces along one category (e.g., race) compared to another category (gender). There is also ERP evidence that neural encoding of similarity of other-race (e.g., Black) faces took place earlier than that of same-race (e.g., Asian or White) faces (Zhou et al., 2020; Zhang et al., 2023), indicating distinct temporal procedures of classification of faces of different subcategories. Despite these findings, it remains unclear whether and how social contexts give priority to categorization of the same set of faces based on one over another level of abstraction of facial attributes.

A recent study suggested that social task demands may modulate the temporal procedures of social categorization of faces by race or gender (Zhang et al., 2023). This study provided both behavioral and ERP evidence that the priority of racial or gender categorization for faces dynamically shifted in response to the social task demands of handling intergroup and cross-sex interactions. Because social task demands may vary across different social contexts, it is possible that changing social contexts may modulate the priority of categorization of faces based on one or another level of abstraction of facial attributes. Specifically, since different social identities may be organized in a hierarchy in which ‘human’ constitutes the superordinate level of a person’s identity and other profiles such as race or nationality constitute the subordinate levels of his/her identity (Stets and Burke, 2000), social categorization of faces may be also organized in a similar hierarchy. When encountering people in a context of non-human animals, initial task demands usually require us to distinguish humans from animals (e.g., talk to humans but not to non-human animals) and such task demands are ubiquitous regardless whom (e.g., Black or White, women or men, adults or children) we encounter. However, the initial task demands may be more nuanced when meeting the same people in the presence of other human beings (e.g., talk to them in Chinese or English and what to talk about). To meet these different task demands, it is thus likely that categorization of faces based on a superordinate level of abstraction (e.g., Species: human or animal) takes place earlier or faster than that based on a subordinate level of abstraction (e.g., Race: Asian or Black).

We tested this hypothesis by collecting behavioral and neuroimaging measures related to facial categorization in three experiments. In Experiment 1 we recorded behavioral responses to Asian faces in two explicit facial classification tasks that required classification of Asian versus dog faces (dog faces provide a species context) or Asian versus Black faces (Black faces provide a race context). According to our hypothesis, we expected faster behavioral responses to Asian faces in the species context compared to the race context. In Experiments 2 and 3 we combined electroencephalography (EEG), magnetoencephalography (MEG), and a repetition suppression (RS) paradigm (Zhou et al., 2020; Zhang and Han, 2021; Zhang et al., 2023) to further examine the neural architecture involved in categorization of Asian faces in the species and race contexts, respectively. The RS paradigm consisted of two conditions in which faces of one social category (e.g., Asian or Black) were presented repeatedly (the repetition condition or Rep-Cond) or faces of two different social categories (e.g., Asian and Black) were displayed alternately in a random order (the alternating condition or Alt-Cond). Decreased neural responses to faces of the same (race or gender) category in the Rep-Cond compared to Alt-Cond have been repeatedly identified (Zhou et al., 2020; Zhang and Han, 2021; Zhang et al., 2023). This RS effect reflected neural encoding of similarity of faces of the same category as a key basis of social categorization of faces. The neural RS effect manifested processes underlying spontaneous categorization of faces because it was observed when participants performed a one-back (i.e., to detect a casual target face that was displayed in two consecutive trials) that prompted individuation rather than categorization. It has been shown that, when Chinese participants viewed Asian and Black faces in Alt-Cond and Rep-Cond, RS occurred to the right fusiform activity in response to Black faces at 140-200 ms but to the activity in the left temporoparietal junction (TPJ) in response to Asian faces at 200-400 ms post-stimulus (Zhou et al., 2020). These results uncovered neural underpinnings spontaneous categorization of Asian faces in a race context but left an open question regarding how the brain classifies Asian faces in a context of non-human animal faces (i.e., the species context).

Early research showed evidence that MEG signals were able to decode human versus animal faces around 120 ms after stimulus onset (Cichy et al., 2014). However, the amplitudes of M170 (a neural response corresponding to the face-specific N170) did not differ significantly between human and monkey faces (Yamada et al., 2025). The results suggested early neural discrimination between human and animal faces but did not examine the neural encoding of similarity of human faces in the context of animal faces. A recent study that combined EEG/MEG and the RS paradigm found that, in the context of human body silhouettes, RS of neural responses to non-human animal body silhouettes occurred in two time windows in the left supramarginal gyrus (around 200 ms) and left frontal cortex (around 400 ms), respectively (Pu and Han, 2025). These findings suggested two neural processes underlying species-based categorization of body silhouettes but not faces. The current work employed EEG/MEG and the RS paradigm to compare the RS effect on neural responses to the same set of Asian faces in the species context and in the race context. If there is a hierarchical organization of spontaneous categorization of faces based on a superordinate (e.g., Species) and a subordinate (e.g., Race) level of abstraction, the RS effect on neural responses to Asian faces as an index of spontaneous categorization of Asian faces would occur earlier in the species context then in the race context. Our study showed consistent behavioral and EEG/MEG results that uncovered context-dependent categorization of faces and its underlying neural architecture.

## Materials and Methods

### Participants

This study recruited 31 participants in Experiment 1 (15 males, mean age ± s.d. = 21.3 ± 2.4 years), 30 participants in Experiment 2 (15 males, 21.3± 2.3 years), and 31 participants for Experiment 3 (15 males, 21.1 ± 2.2 years). All participants were Chinese college students. All were right-handed, had normal or corrected-to-normal vision and reported no history of neurological or psychiatric diagnoses. All participants provided written informed consent after the experimental procedure had been fully explained. Participants were reminded of their right to withdraw at any time during the study and were paid for their participation. This study was approved by the local ethics committee at the School of Psychological and Cognitive Sciences, Peking University.

### Stimuli

The stimuli used in this study consisted of three sets of (Asian, black and dog) faces. Sixteen Asian (8 males) and 16 black (8 males) faces with neutral expressions were adopted from the RADIATE face stimulus set (Conley et al., 2018). Sixteen photos of frontal views of dog faces with neutral expressions were collected from internet collections. All faces were cropped to remove surrounding hair and to ensure a similar face configure (a near oval shape with two eyes above a nose and a mouth). All photos were transformed into gray-scale images with 200 × 250 pixels. The luminance levels of the three sets of stimuli were adjusted and matched. Each face subtended a visual angle of 4.0° × 5.0° at a view distance of 80 cm in Experiment 1 and the behavioral classification task in Experiment 3. Each face subtended a visual angle of 4.0° × 5.0° at a view distance of 80 cm in the EEG experiment. Each face subtended a visual angle of 6.9 × 9.9° at a view distance of 75 cm in the MEG experiment.

### Face classification task

A face classification task was conducted to estimate behavioral performances during classification of the Asian faces in different contexts in Experiments 1 and 3. This task was also conducted in Experiment 3 after MEG recording. There were 4 blocks of 32 trials in which faces of two categories were presented in a random order. Asian faces were displayed alternately with Black faces (the race context) in 2 blocks of trials and with dog faces (the species context) in 2 blocks of trials. Each face was presented in the center of a grey background for 200ms and was followed by a fixation cross for 1200ms. The participants were instructed to sort each face into one of the two categories by pressing one of two keys as quickly and accurately as possible. Data analyses focused on the difference between behavioral responses to Asian faces in the race context and species contexts. Responses shorter than 200ms or responses longer than 1200 ms were excluded from data analyses, resulting in elimination of <1% of trials.

### Procedures of EEG and MEG experiments

During EEG recording in Experiment 2 each trial consisted of a face in the center of a grey background with a duration of 200 ms, which was followed by a fixation cross with a duration varying randomly from 250 to 550 ms. Participants were asked to respond to the immediate repetition of the same face in two successive trials by pressing a button (the one-back task). Each participant completed two EEG recording sessions in which Asian faces were presented in the race context or the species context, respectively. The order of the two sessions was counterbalanced across participants. There were three runs for each EEG recording session. Each run consisted of 8 blocks (4 blocks in the Rep-Cond and 4 blocks in the Alt-Cond) of 18 trials (16 non-target faces and 2 target faces in each block). There was a 5-second break between the two consecutive blocks in which the number 5 was display at the fixation position and flashed with a 1-s step and decreased to 1. Faces were presented in a random order in each block.

The stimuli and procedure during MEG recording in Experiment 3 were the same as those in Experiment 2 except for the following. There were four runs for each MEG recording session and 8 blocks for each run. After MEG recording participants completed the face classification task.

### EEG data acquisition and analysis

EEG signals were continuously recorded using Brain Vision Recorder (Brain Products, GmbH) with 64 Ag-AgCl scalp electrodes placed according to the International 10-20 system and referenced online to the FCz electrode. AFz was used as the ground electrode. Impedances of individual electrodes were kept below 5 kΩ. Eye blinks and vertical eye movements were monitored with electrodes located below the right eye. The EEG was amplified (band pass 0.1–1000 Hz) and digitized at a sampling rate of 500 Hz.

EEG data were analyzed with Brain Vision Analyzer 2.0 (Brain Products, GmbH). EEG data were re-referenced offline to the average of the left and right mastoid electrodes (TP9/TP10) and then filtered with low-pass filter at 40 Hz. Eye-movement artifacts were corrected by conducting the semiautomatic independent component analysis (ICA) application in Analyzer 2.0. The ERPs in each condition were averaged separately offline with an epoch beginning 200 ms before stimulus onset and continuing for 650 ms. Only EEG signals to non-target Asian faces (96 trials in each condition) were included for analyses. Trials contaminated by noise exceeding ±100 μV at any electrode were excluded from average. This resulted in 85 ± 4 trials accepted per condition in Experiment 2. The baseline for ERP measurements was the mean voltage of a 200 ms prestimulus interval. The latency of each ERP component was measured relative to the stimulus onset.

Event-related potentials (ERPs) to non-target Asian faces were characterized by a negative activity at 120-140 ms (N1) followed by a positive activity at 156-216 ms (P2) and a negative activity at 220-420 ms (N2) over the frontal/central electrodes. The mean values of the amplitudes of the N1, P2 and N2 components were calculated at frontal (Fz, FCZ, F3, F4, FC3 and FC4) and central (Cz, C3 and C4) electrodes. To avoid any bias due to the selection of time windows for ERP amplitude measurements, we averaged the ERPs across different conditions and then used the timing and scalp distribution of these averaged ERPs to define the time windows for measurements of mean amplitudes of ERP components. The peak latency ± full width at half maximum was set up as the time window for measuring the mean P2 amplitude. The mean amplitudes were then subject to repeated-measures analysis of variance (ANOVA) with condition (Rep-Cond versus Alt-Cond) and context (Race versus Species contexts) as within-subject variables. RS effects were defined as decreased amplitudes to non-target Asian faces in the Rep-Cond compared to Alt-Cond. ERPs to non-target Asian faces also elicited a negative going wave N170 at 160–220 ms over the bilateral occipito-temporal electrodes (P7 and P8). Similarly, the mean N170 amplitudes were subjected to ANOVAs with condition (Rep-Cond versus Alt-Cond) and context (Race versus Species contexts) as within-subject variables.

### MEG data acquisition and analyses

#### MEG and structural MRI data acquisition

Cortical neuromagnetic activity was recorded using a whole-head MEG system with 102 magnetometers and 204 planar gradiometers (Elekta Neuromag TRIUX) in a magnetically shielded room. The MEG signals were sampled at 1 kHz with an online 0.1–330 Hz band-pass filter. Structural MRI of all the subjects’ heads were collected using a Siemens Trio 3.0 T MR scanner with a 12-channel phase-array head coil at the Center for MRI Research, Peking University. A high-resolution anatomical T1 - weighted image was acquired for each participant (256×256mm matrix, 192 slices, 1×1×1 mm^3^ voxel size; TR = 2530ms, TE=2.98ms, inversion time (TI)=1100ms, FOV = 25.6×25.6 cm, FA = 7°, scanning order: interleaved). To co-register the MEG data with MRI coordinates, 3 anatomical landmarks (nasion, left, and right pre-auricular points), 4 HPI coils and at least 200 points on the scalp were digitized using the Probe Position Identification system (Polhemus, VT, USA). During the MEG scan, the head coil positions were repeatedly recorded by electromagnetic induction at the beginning and end of each block. A Maxfilter software (Elekta-Neuromag, Helsinki, Finland) with temporal signal space separation (tsss) was first used to remove the external interference to the raw MEG data. For each condition, the head positions of each pair of blocks were coregistered in reference to the position of the first block using Maxmove (a sub-component of Maxfilter software) before combining the MEG data. *Sensor-space whole-brain analysis*.

The offline analysis of MEG data was performed using Brainstorm package (Tadel et al., 2011). For time-course analysis, a low-pass filter of 40 Hz was applied. Eye blink artifacts were attenuated with signal space projection (SSP) by visually inspecting and removing the corresponding SSP component. The data were then epoched in accordance with stimulus trigger codes. Each epoch started 200 ms before face onset and continued for 650 ms. Rejection parameters for further data quality inspection were set at 3,500 fT/cm for gradiometers, 3,500 fT for magnetometers, which resulting in the inclusion of 104±4 trials in each condition. The response to each face was baseline-corrected based on the 200-ms period preceding the face onset for each sensor.

A whole-brain cluster-based permutation t-test (Maris and Oostenveld, 2007) was used to detect significant RS effects for Asian faces on magnetometer signals and averaged norm for the planar gradiometer signals. For each context, a t-value was computed between MEG signals to Asian faces in the Rep-Cond and the Alt-Cond. Adjacent points in time and space exceeding a predefined threshold (*P* < 0.05, two-tailed) were grouped into one or multiple clusters. The summed cluster t-values were compared against a permutation distribution that was generated by randomly reassigning condition membership for each participant (10,000 iterations) and computing the maximum cluster mass for each iteration. This approach reliably controls for multiple comparisons at the cluster level. The permutation tests were performed at 0-450 ms because it takes at least 450 ms before the presentation of the next face stimulus. The RS effect at cluster level was defined as the absolute value of the Alt-Cond minus the Rep-Cond when the original value of the two conditions had the same sign. The significant clusters were defined using a cluster-level threshold of P < 0.05 with corrections for multiple comparisons. Because one magnetometer channel (MEG2031) was broken, magnetometer data from 101 channels were subject to data analyses.

#### Source-space whole brain analysis

An anatomical T1 scan was acquired for each participant for source constructions of MEG signals. T1 images were obtained using a Siemens Trio 3.0 T MR scanner with a 12-channel phase-array head coil at the Center for MRI Research, Peking University. Segmentation of T1 images was conducted using automated algorithms provided in the software package FreeSurfer (Blanco-Elorrieta et al., 2018) (http://surfer.nmr.mgh.harvard.edu/).

Source localization and surface visualization were performed using the toolbox BrainStorm (Tadel et al., 2011). After co-registration between the individual anatomy and MEG sensors, cortical currents were estimated using a distributed model consisting of 15,002 current dipoles from the combined time series of magnetometer and gradiometer signals using a linear inverse estimator (weighted minimum-norm current estimate, signal-to-noise ratio of 3, depth weighting of 0.5) separately for each condition and for each participant in a single-sphere head model. Dipole orientations were constrained with help of individual high-resolution cortical surface reconstructions. Noise covariance matrix was acquired from 2 min of empty-room MEG recordings collected daily before the experiment. For the group analysis, individual source-space data were projected to a standard brain model (Colin27, 15,002 vertices) using BrainStrorm (Tadel et al., 2011). For each of the 15,002 vertices, source activations were obtained by standardizing those values to pre-stimulus intervals of 200 ms (subtracting the mean and dividing by the standard deviation of the baseline) and computing the absolute value of each dipole. We also applied spatial smoothing using a 3-mm FWHM Gaussian kernel. For the purposes of group statistical analysis in the source space, the activity of each vertex over the time windows, in which significant activations were observed in the sensor space, was averaged. Whole-brain analyses of significant RS effects (that is, decreased neural responses to Asian faces in the Rep-Cond versus the Alt-Cond) in source-space similarly used the cluster-based permutation t-test performed on 15,002 vertices of the cortical surface model (using a predefined threshold of *P* < 0.01, one-tailed, 10,000 iterations). One-tailed tests were implemented here to examine decreased neural responses to Asian faces in the Rep-Cond versus the Alt-Cond. The significant clusters were defined using a cluster-level threshold of *P* < 0.05 with corrections for multiple comparisons.

#### Source-space spatiotemporal point of interests (STOI) analysis

We further performed STOI analyses to examine the RS effects on source-space MEG signals to Asian faces in different contexts. Spatiotemporal points were defined based on 50 vertices in each ROI in which the mean brain activity in a specific time window showed reliable RS effects on source-space MEG signals (at 140–200 ms in the right fusiform gyrus, RFG, peak MNI coordinates: x, y, z = 47, -52, -24; at 317-413 ms in the left inferior parietal cortex (LIP), supramarginal gyrus (LSMG), and left temporoparietal junction (LTPJ), peak MNI coordinates: x, y, z = -36, -86, 39). The mean value of each STOI were then subject to repeated-measures ANOVA with condition (Rep-Cond versus Alt-Cond) and context (Race versus Species contexts) as within-subject variables. To further assess the null hypothesis in early time stage in race context, we conducted Bayes factor analyses for each RS effect. We calculated the Bayes factor in the program R v.3.5.1 (www.r-project.org) using the function ttestBF from the package BayesFactor (Morey et al., 2015). A Bayes factor indicates how much more likely each alternative model is supported compared with the null hypothesis.

## Results

### Behavioral evidence for context-dependent hierarchical categorization of faces

In Experiment 1 we tested our hypothesis by examining contextual effects on behavioral responses in the face classification task. Reaction times and response accuracies to the same set of Asian faces were recorded from Chinese adults (*N* = 31) in the race-context (Asian and Black faces were presented in a random order) and the species -context (Asian and dog faces were presented in a random order), respectively. The participants had to sort each face into one of the two categories (Asian vs. Black in the race context and human vs. dog in the species context) as fast and accurately as possible. Response efficiencies pertaining to Asian faces in each task were calculated by dividing the mean reaction times by the mean response accuracy. A smaller performance efficiency indicates faster behavioral classification of faces. If categorization of faces based on a superordinate level of abstraction (e.g., Species: human or animal) takes place earlier or faster than that based on a subordinate level of abstraction (e.g., Race: Asian or Black), we would expect smaller response efficiencies related to Asian faces in the species context than in the race context.

Indeed, a paired-*t* test revealed that the performance efficiency was significantly smaller in response to Asian faces in the species-based compared to race-based classification task (*t*(30) = 6.22, p < 0.001, Cohen’s d = 1.12, mean difference = 91.9, 95% CIs = [61.7, 122.1], Fig.1; see Table S1 for details of accuracies and reaction times). A post-hoc power analysis using G*Power (Faul et al., 2007) showed that the sample size in Experiment 1 had a power over 0.99 to reveal a reliable difference in response efficiencies between the two conditions. Because the same set of Asian faces were used in the two classification tasks and the order of the two classification tasks were counterbalanced across the participants, the results observed here cannot be explained by any effects of perceptual features of the stimuli or response habituation but manifested influences of the contexts. The results favor the hypothesis of hierarchical face categorization that occurs earlier or faster based on a superordinate (i.e., Species) compared to a subordinate (i.e., Race) level of abstraction of facial attributes.

**Figure 1.**
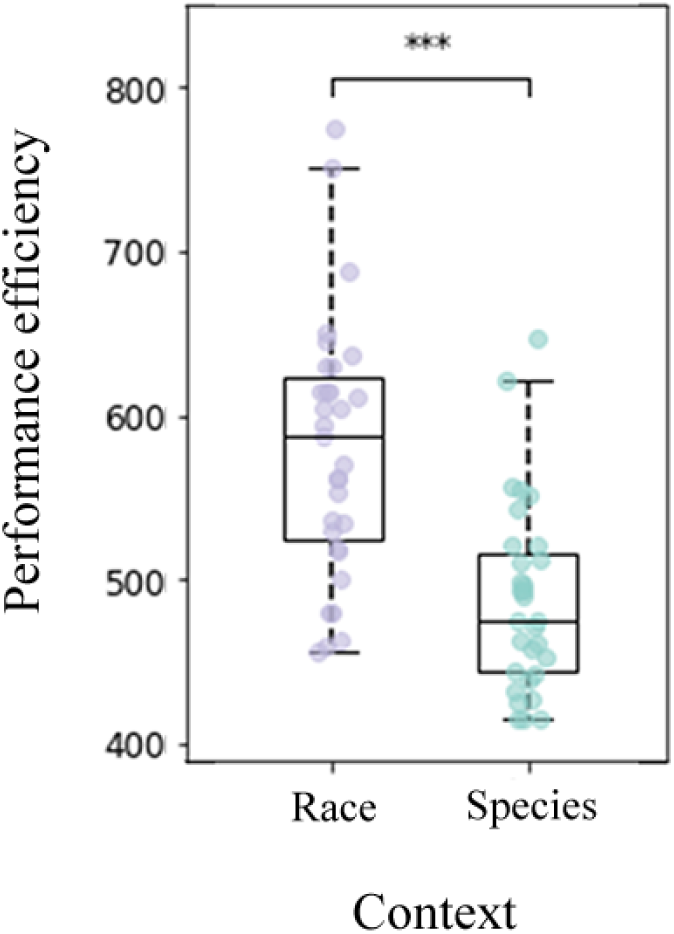
Behavioral results in Experiment 1. Performance efficiency related to Asian faces was smaller (indicating earlier or faster responses) in the species-based relative to race-based contexts.

### EEG evidence for context-dependent hierarchical categorization of faces

The decreased performance efficiency related to Asian faces in the species than race context in Experiment 1 reflected the final outcomes of the contextual effect but did not disentangle the exact temporal procedure of categorization of faces in the two contexts. In Experiment 2 we employed EEG and the RS paradigm (Zhou et al., 2020; Zhang et al., 2023) to address this issue by examining neural encoding of similarity of different Asian faces in the species context and race context, respectively. In the Rep-Cond different faces of one category (i.e., Asian, Black, or dog) were presented in a random order. In the Alt-Cond different faces of two categories (i.e., Asian and Black in the race context, Asian and dog in the species context) were presented in a random order. The participants responded to a casual target face that was displayed in two consecutive trials by a button press (i.e., the one-back task). EEG signals to faces in the Alt-Cond and Rep-Cond were collected from an independent sample of Chinese adults (*N* = 30). We focused on the RS effects on neural responses to non-target Asian faces (i.e., deceased neural activities in response to Asian faces in the Rep-Cond than in the Alt-Cond) in the race and species context, respectively. The RS effect allowed us to examine the temporal procedure of spontaneous neural encoding of similarity of Asian faces in the two different contexts. According to the hypothesis of hierarchical categorization of face, we predicted that the RS effect on neural responses to Asian faces would occur earlier in the species context than in the race context

Participants detected targets with high response accuracies in the one-back task (>88%, see Table S2 for details). The repeated-measures analysis of variance (ANOVA) of reaction times and response accuracies related to Asian faces with Condition (Alt-Cond versus Rep-Cond) × Context (Race versus Species) as within-subject variables did not show any significant effect (see Table S2 for statistical details), suggesting comparable attentional demand and task difficulty between detections of Asian targets in the species and race contexts. ERPs to non-target Asian faces were characterized by a negative activity at 120-140 ms (N1) followed by a positive activity at 156-216 ms (P2) and a negative activity at 220-420 ms (N2) over the frontal/central electrodes (Fig. 2A & 2B), and negative activity at 160-220 ms (N170) over the lateral occipitotemporal electrodes (see Fig. S1). The mean amplitude of each component at these electrodes was subject to ANOVAs with Condition (Alt-Cond versus Rep-Cond) × Context (Race versus Species) as within-subject variables to examine the RS effects on neural responses to non-target Asian faces in different contexts (see Table S3 for statistical details).

**Figure 2:**
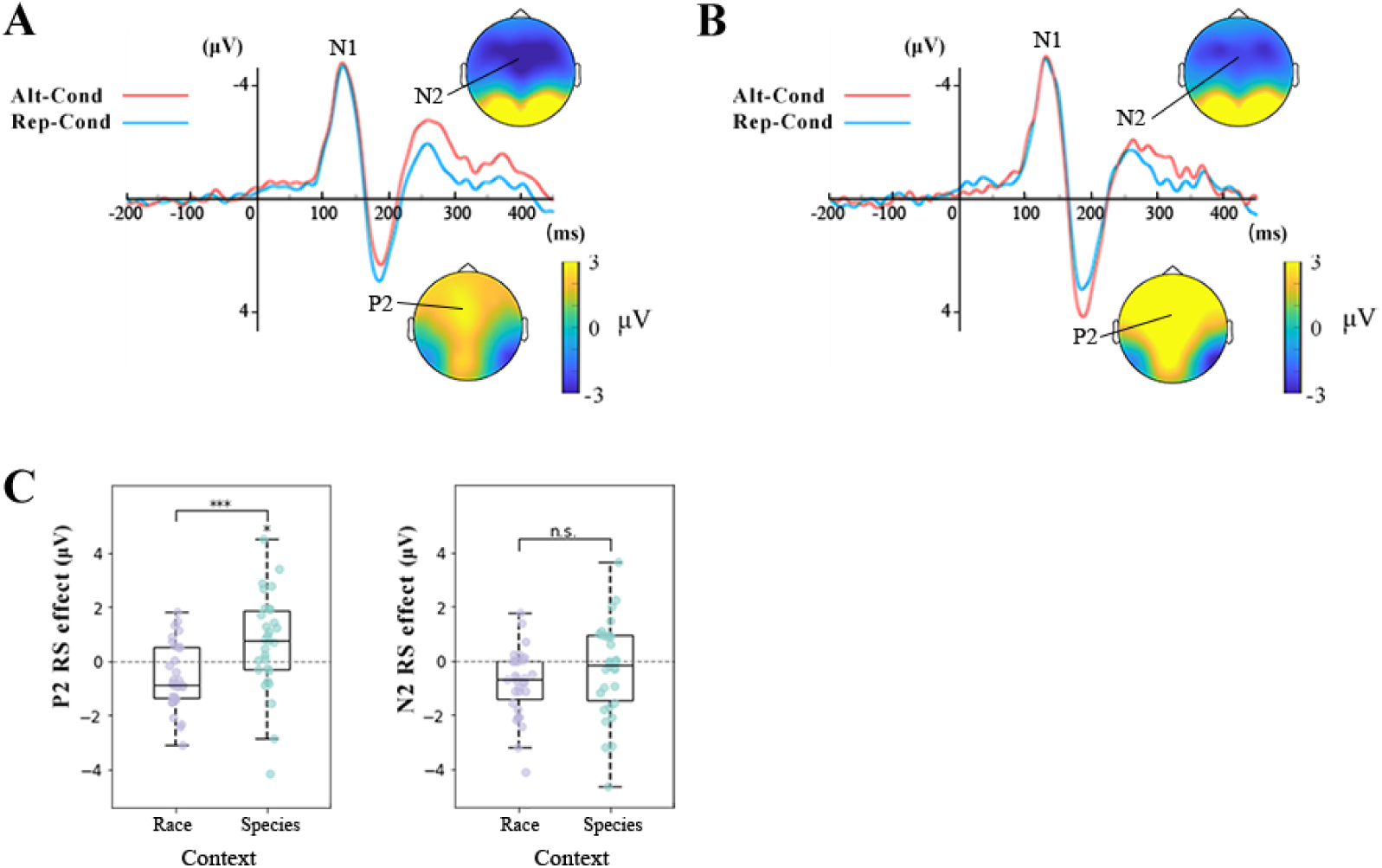
ERP results in Experiment 2. (A) Illustrations of ERPs to non-target Asian faces at the frontal/central electrodes in the race context. (B) Illustrations of ERPs to non-target Asian faces at the frontal/central electrodes in the species context. (C) The mean RS effects related to non-target Asian faces in the P2 and N2 windows in the two contexts.

ANOVAs of the P2 amplitudes to Asian faces showed a significant main effect of Context (*F*(1,29)=11.91, *p*=0.002, *ƞ^2^_p_*=0.291, mean difference=0.840, 90% Cis =[0.291,0.939]), suggesting larger P2 amplitudes to Asian faces in the species context than in the race context. Importantly, there was a significant interaction of Condition × Context (*F*(1,29)=13.372, *p*=0.001, *ƞ^2^_p_*=0.316, 90% CIs =[0.324,0.979]. Simple effect analyses revealed reliable RS effects on the P2 amplitudes to Asian faces in the species context (*p*=.043, mean difference=.698), but not in the race context (a repetition enhancement effect emerged, *p*=0.021, mean difference=-.553, (Fig. 2C). These results provide evidence that the RS effect on neural responses to Asian faces occurred in the P2 time window in the species context but not in the race context. ANOVAs of the N2 amplitudes to Asian faces showed a significant effect of Condition (*F*(1,29)=6.04, *p*=.02, *ƞ^2^_p_*=0.172, 90% CIs =[0.125,0.748]), due to larger N2 amplitudes in Alt-Cond than in Rep-Cond. However, this effect did not differ between the species and race contexts (*F*(1,29)=20553, *p*=.122, *ƞ^2^_p_*=0.080, 90% Cis =[0, 0.585]). These results are consistent with the previous findings (Zhou et al., 2020; Zhang et al., 2023) and indicates neural encoding of attributes of Asian faces in the N2 window in both the species and race contexts. ANOVAs of the N1 and N170 amplitudes to Asian faces did not show any significant effect (*ps* > 0.22).

Together, the ERP results provided ERP evidence for the RS effects on neural responses to Asian faces. More importantly, the ERP results unraveled distinct temporal procedures of the neural encoding of attributes of Asian faces in the species and race contexts. The earlier neural encoding of attributes of Asian faces in the species compared to race contexts provide electrophysiological evidence for a hierarchy of spontaneous neural categorization of faces. In addition, these ERP results suggested two stages of categorization of Asian faces in the species context in the P2 and N2 time windows, respectively. By contrast, categorization of Asian faces in the race context was evidence only in the late N2 time window.

### Neural circuits underlying the hierarchical categorization of faces

To further investigate the neural circuits involved in hierarchical categorization of faces, in Experiment 3, we recorded 306-channel, whole-head anatomically constrained MEG signals from an independent sample of Chinese adults (*N*=31). The stimuli and procedures were the same as those used in Experiment 2 except that there were 128 trials in each condition (see Methods for details, and see Table S4 for behavioral results). The participants also completed the same face classification task used in Experiment1 after MEG recording (see Table S5 for details of the results). We conducted source estimation of the RS effects on neural responses to Asian faces in the species and race contexts and examined possible relationships between the neural RS effects and behavioral performances in the face classification task.

We first conducted cluster-based permutation t-tests to examine the RS effects on sensor-space MEG signals in response to Asian faces in the species context. A significant RS effect was identified using a predefined threshold of P < 0.05, two-tailed, 10,000 iterations, and a cluster-level threshold of P < 0.05 with corrections for multiple comparisons. The results showed significant RS effects on sensor-space MEG signals to Asian faces in the species context at the right occipito-temporal regions (140–200 ms, cluster *p* = 0.004, magnetometer signals; cluster *p* = 0.007, gradiometer signals) and the left occipito-parietal region (317-413 ms, cluster *p* = 0.005, gradiometer signals, Fig. 3A & 3B). These results replicated our ERP results in Experiment 2 and provide MEG evidence for early neural encoding of similarity of Asian faces in the species context.

**Figure 3:**
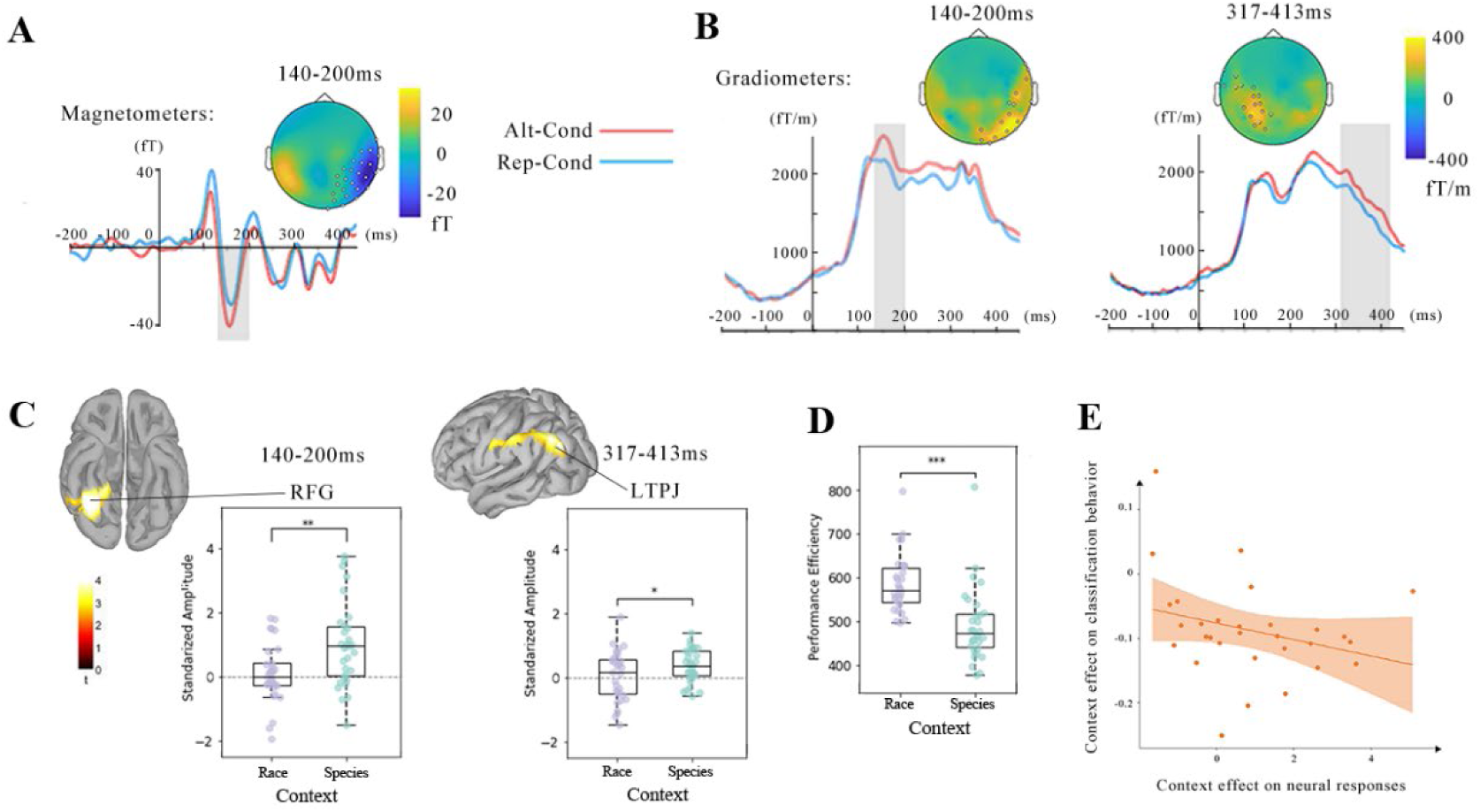
MEG results in Experiment 3. (A) and (B) RS effects on sensor-space MEG signals in response to Asian faces in the species context. (C) RS of source-space MEG signals to Asian faces in the species context in the RFG and left TPJ. The RS effect on responses to Asian faces was significantly greater in the species context than in the race context. (D) Better behavioral performances (i.e., smaller performance efficiency) related to Asian faces in the classification task in the species context than in the race context. (E) Greater contextual effects on RS of neural responses to Asian faces in the right fusiform gyrus were associated with greater contextual effects on the response efficiencies related to Asian faces in the classification task. The x-axis represents the difference in the RS effect (species context - race context) in the right fusiform gyrus STOI (140-200ms). The y-axis represents the difference in performance efficiency (species context - race context) in the classification task. FG = right fusiform gyrus; TPJ = temporoparietal junction.

Next, we conducted whole-brain source-level analyses to estimate sources of the RS effects on neural activities in response to Asian faces in the species context. A significant RS effect in a brain region was identified by conducting one-tailed tests to examine decreased neural responses to Asian faces in the Rep-Cond compared to the Alt-Cond (using a predefined threshold of *P*<0.01, one-tailed, 10,000 iterations, and a cluster-level threshold of *P*<0.05). The results yielded two clusters showing significant RS effects on neural responses to Asian faces in the right fusiform gyrus (RFG) at 140–200 ms (peak MNI coordinates: x, y, z = 47, -52, -24; cluster *p* = 0.008) and in the left inferior parietal cortex (LIP)/supramarginal gyrus (LSMG)/temporoparietal junction (LTPJ) at 317-413 ms (peak MNI coordinates: x, y, z = -36, -86, 39; cluster *p* = 0.022, Fig. 3C).

Because the whole-brain sensor-level and source-level analyses did not find significant RS effect on neural responses to Asian faces in the race context, we performed spatiotemporal points of interests (STOI) analyses to compared the RS effects on source-space signals in the species and race contexts. Spatiotemporal points included 50 vertexes around the peak points in the RFG (47/-52/-24) and the LSMG/LTPJ (-36/-86/39). Averaged neural activities of each STOI among vertices and across time points were extracted and subject to ANOVAs with Condition (Rep-Cond versus Alt-Cond) and Context (Race versus Species contexts) as within-subjects variables. The results showed significant two-way interactions on MEG signals in both the RFG (*F*(1,30) = 8.52, *P* = 0.007, *ƞ^2^_p_* = 0.221, 90% CIs = [0.201,0.823]) and LIP/LSMG/LTPJ (*F*(1,30) = 5.55, *P* = 0.025, *ƞ^2^_p_* = 0.156, 90% CIs = [0.105,0.717]), indicating stronger RS effects on neural responses to Asian faces in the two regions in the species context than in the race context (Fig. 3C). We further conducted Bayes factor analyses to assess the evidence for RS effect to Asian faces in the race context in each STOI. The results showed medium evidence for RS effects only on the late LIP/LSMG/LTPJ signals (BF_10_ = 4.326), consistent with the previous finding (Zhou et al., 2020). These results revealed RS of both early neural responses to Asian faces in the RFG and later neural response to Asian faces in the LIP/LSMG/LTPJ in the species context but of only the later neural response to Asian faces in the LIP/LSMG/LTPJ in the race context.

Similarly, we found a significantly smaller performance efficiency in response to Asian faces in the species-based compared to race-based classification task in Experiment 3 (t(29) = 6.31, p < 0.001, Cohen’s d = 1.15, mean difference = 87.5, 95% CIs = [59.1, 115.8], see Fig. 3D, Table S5). For each participant, we calculated the difference in performance efficiency between two contexts and defined it as the contextual effect on classification behavioral performances. We also extracted the mean RS effects in the right fusiform gyrus STOI (140-200ms) in the species and race contexts and calculated the difference to assess the contextual effect on the neural categorization of Asian faces. A Spearman correlation analysis revealed a negative correlation between the contextual effects on classification behavioral performances and neural categorization of Asian faces (Spearman rho = -0.36, p = 0.495, Fig. 3E), suggesting that a stronger contextual effect on neural categorization of Asian faces was associated with a greater contextual effect on behavioral classification of Asian faces.

## Discussion

The present study investigated the impacts of social contexts on behavioral and neural responses to faces to test a possible hierarchical structure of social categorization of faces based on a superordinate (species) and a subordinate (race) level of abstraction of facial attributes. Experiment 1 showed that the species context compared to the race context accelerated behavioral responses to Asian faces in the explicit facial classification task, providing behavioral evidence for faster categorization of Asian faces in the species context compared to the race context. Experiment 2 found that neural encoding of similarity of Asian faces, which was indexed by the RS effects on ERP amplitudes to Asian faces and manifested a key neural underpinning of spontaneous categorization of faces, occurred in both the P2 time window (156-216 ms) and the N2 time window (220-420 ms) in the species context but only in the latter time window in the race context. These ERP results provided EEG evidence for earlier spontaneous categorization of faces in the species context than in the race context. Our MEG results in Experiment 3 showed that the RFG and LIP/LSMG/LTPJ were engaged in encoding of similarity of Asian faces in the two successive time windows in the species context, whereas only the LIP/LSMG/LTPJ activity in the later time window was engaged in encoding of similarity of Asian faces in the race context. In addition, the contextual effects on behavioral responses to classification of Asian faces were associated with those on the fusiform activity underlying encoding of similarity of Asian faces. Together, these behavioral and EEG/MEG findings unveil a temporal/spatial hierarchical structure of categorization of faces based on a superordinate (Species) and subordinate (Race) levels of abstraction of social attributes.

Context-dependent categorization of human faces underscores the adaptive function of face perception in diverse social contexts. Previous research has suggested effects of contextual information on facial categorization across various dimensions. For example, visual contexts of American-typed or Chinese-typed scenes modulated explicit racial categorization by biasing perception of racially ambiguous faces toward categories congruent with the scene context (Freeman et al., 2013, 2015). Socioeconomic contexts implying poverty (vs. wealth) led to a greater tendency for the middle-class White participants to classify same-race faces as out-group members (Shriver et al., 2008). Contextual features such as hairstyles that act as racial markers also shape the explicit categorization of ambiguous faces into a racial category (MacLin and Malpass, 2001, 2003). Despite these findings, there has been a lack of in-depth discussion on how social categorization of faces is organized in terms of the contexts that may give prominence to different levels of facial attributes. Moreover, prior studies typically employed tasks that prompted participants to explicitly classify faces based on different levels of abstraction or along different dimensions (Bowers and Jones, 2008; Farzmahdi et al., 2021). In real-life situations, however, social categorization of faces usually takes place spontaneously rather than in response to explicit instructions. Our EEG/MEG findings indicate a hierarchy of spontaneous categorization of faces that occurs early in a context that made salience the superordinate (Species) level of abstraction of faces but delayed in the context that highlighted the subordinate (Race) level of abstraction of faces. In addition, we showed that categorization of faces based on species may engage successive activities in two neural circuits in the right fusiform gyrus and the LIP/LSMG/LTPJ whereas categorization of faces based on race was linked only to the late activity in the LIP/LSMG/LTPJ.

The fusiform gyrus is well known for its functional role in face perception. The fusiform activity responded more strongly to human faces relative to objects and bodies (Kanwisher et al., 1997), indicating early neural representations of inter-category (faces versus non-faces) differences. The fusiform activity also showed habituation in response to repetition of the same face (Gauthier et al., 2000), suggesting its key role in neural representation of individuals’ identities. However, fMRI studies found that neural activities in the fusiform may distinguish between faces of two racial categories (Golby et al., 2001; Lovén et al., 2014). Recent research using the RS paradigm also found evidence for RS of the fusiform activity in response to other-race (Black) faces but not to own-race (Asian) faces (Zhou et al., 2020), suggesting that the fusiform is also involved in categorical representations of Black (but not Asian) faces. The findings of the current work suggest that the fusiform may contribute to the clustered representations of Asian faces when the context gave prominence to the facial attributes of humanness. While the previous finding of greater fusiform responses to human compared to animal faces (Kanwisher et al., 1999) manifested distinction between representations of human and animal faces as two different categories, our findings highlighted the effect of the species context on facilitation of the clustered representations of human faces in the fusiform.

Research on monkeys has revealed evidence for the critical role of the intraparietal activity in transforming of visual feature selectivity to abstract categorical representations of stimuli for appropriate behavioral responses (Freedman and Assad, 2006). Recent MEG findings found that the LIP/LSMG supports clustered representations of animal body silhouettes in the context of human body silhouettes (Pu and Han, 2024). The TPJ responses more strongly to social category-related behavioral information than individual-related trait-related information (Van der Cruyssen et al., 2015) and is involved in social categorization by narrative roles (Ron et al., 2022). Our previous research (Zhou et al., 2020) and the current work showed evidence that the LTPJ is involved in encoding the similarity of Asian faces in both the race and species contexts, indicating that the LTPJ may support categorization of the same set of faces regardless the context changes (i.e., other-race faces or non-human animal faces). These findings together highlight a domain-general function of the neural circuits in the LIP/LSMG/LTPJ in categorical representations of stimuli independent of their perceptual properties and contexts. It is likely that categorical representations at this stage may reflect the conceptual similarity of different individuals rather than perceptual similarity of stimuli.

Our findings have several theoretical implications for understanding social categorization and the underlying neural underpinnings. First, our findings provide converging behavioral and EEG/MEG evidence for a hierarchical organization of categorization of faces based on the superordinate and subordinate levels of facial attributes and revealed the temporal/spatial characteristics of the neural processes involved. Our findings inspire new research questions regarding whether social categorization based on race, gender, and age is organized hierarchically in terms of the temporal/spatial characteristics of underlying neural activities. For example, racial categorization of faces occurred regardless whether faces of one gender or two genders were perceived (Zhou et al., 2020), suggesting dominance of categorization of faces by race relative to that by gender. Such a hierarchical organization of social categorization may also exist in other domains of social information such as body silhouettes (Pu and Han, 2024) but requires future investigation. Second, our findings of a hierarchical organization of categorization of faces fits the social task demand theory of social categorization (Zhang et al., 2023) by showing that various social contexts that drive different social tasks may accelerate or delay categorization of the same set of faces and providing further evidence that social categorization of faces is flexible and adaptive to optimize social interactions. Third, our findings demonstrate that, rather being rigidly determined by perceived stimuli, clustered neural representations of faces is flexible and context dependent. In other words, the neural mechanisms underlying social categorization of faces along different facial attributes may be modulated by social contexts that determine the social task at hand. Finally, our EEG/MEG findings underscore the influence of social contexts on social categorization of faces while the participants performed the one-back task that required individuation rather than categorization of faces. Therefore, social contexts may exert influences on spontaneous (rather than task-driven) categorization of faces that often occurs in real-life situations to facilitate social interactions and manage social relationships effectively. Future research should clarify whether the hierarchical organization of social categorization of faces observed in the current work can be observed during explicit task-driven social categorization of faces

In conclusion, our behavioral and EEG/MEG results showed consistent evidence for the impact of social contexts on spontaneous social categorization of faces. Our findings revealed earlier categorization of faces based on the abstraction of superordinate-level (species) than subordinate-level (race) attributes of faces. The early stage of social categorization of faces was localized to the right fusiform gyrus in which the activity involved in social categorization of faces was enhanced by the species-compared to race-context. By contrast, the late stage of social categorization of faces in the LIP/LSMG/LTPJ was relatively less sensitive to the social contexts. These findings suggest a context-dependent hierarchical structure of social categorization of faces which is consistent with the social task demand framework of social categorization of faces (Zhang et al., 2023). These findings improve our comprehension of social categorization of faces by highlight a hierarchical structure of social categorization of faces as a basis of adaptive social interactions.

## Author contributions

Xuena Wang (Data curation, Formal analysis, Investigation, Methodology, Validation, Visualization, Writing—original draft); Shihui Han (Conceptualization, Data curation, Formal analysis, Funding acquisition, Investigation, Methodology, Project administration, Resources, Software, Supervision, Validation, Visualization, Writing—original draft).

## Data and code availability

The data and the codes used to generate the figures in this study are openly available to the public at https://osf.io/ahce7/?view_only=856426efb5a7481d8ef65815e8166317

## Declaration of generative AI and AI-assisted technologies in the writing process

During the writing process of this work the author(s) did not use any generative AI and AI-assisted technologies.

## Supplementary Results

**Figure S1.**
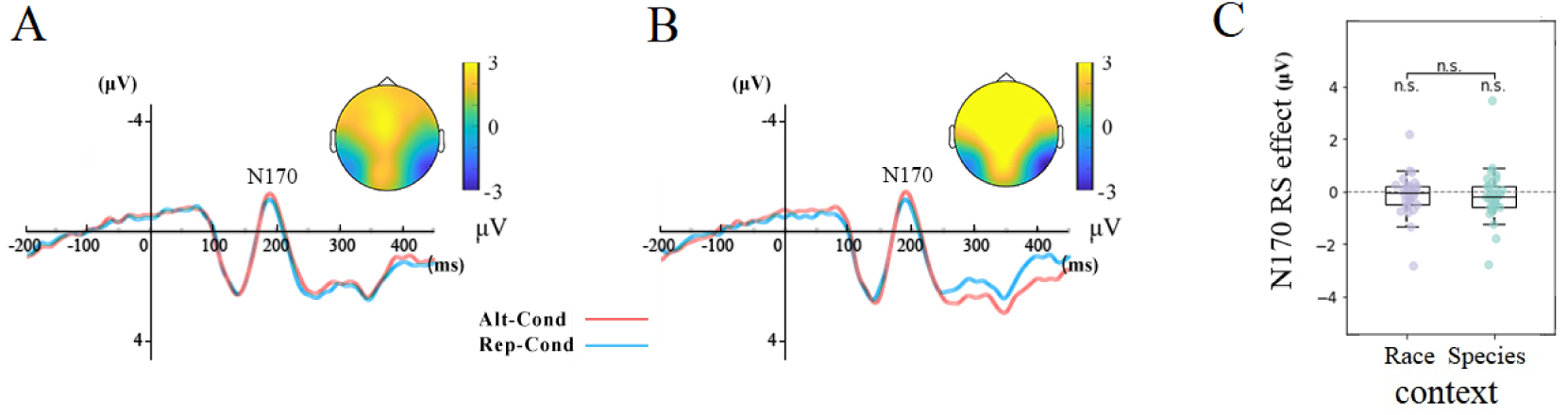
Experimental procedure and results in Experiment 2. (A) Illustrations of presentations of faces in the Alt-Cond and Rep-Cond in the race context and ERPs to non-target Asian faces over the lateral occipitotemporal electrodes. (B) Illustrations of presentations of faces in the Alt-Cond and Rep-Cond in the species context and ERPs to non-target Asian faces over the lateral occipitotemporal electrodes. (C) The RS effects related to non-target Asian faces in the N170 time windows in two contexts. (n.s.: no significance)

**Table S1:**
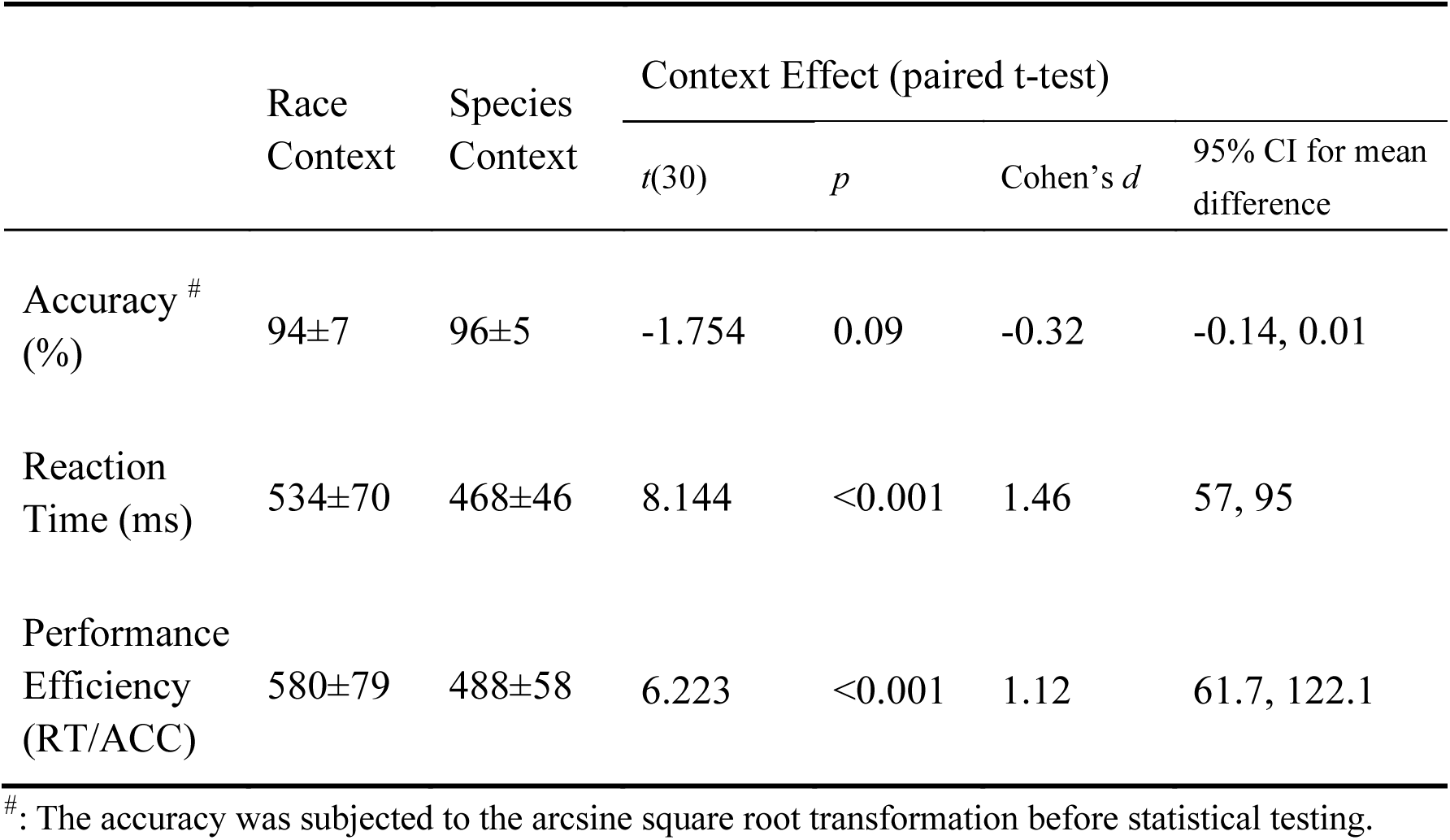
Categorization behavior performance in Experiment 1.

**Table S2:** Behavior performance in the one-back task in Experiment 2.

|  |  | Accuracy (%) <sup>#</sup> | Reaction Time (ms) |
| --- | --- | --- | --- |
| Race context |  |  |  |
|  | Alt-Cond | 90±8.6 | 563±79.0 |
|  | Rep-Cond | 92±9.8 | 555±60.5 |
| Species context |  |  |  |
|  | Alt-Cond | 88±14.8 | 562±54.3 |
|  | Rep-Cond | 90±11.2 | 544±58.8 |
| ANOVA |  |  |  |
| Condition | *F* | 3.243 | 2.794 |
|  | *p* | 0.082 | 0.105 |
| | $\eta^2_p$ | 0.101 | 0.088 |
|  | 90% CI <sup>##</sup> | 0, 0.624 | 0, 0.600 |
| Context | *F* | 0.543 | 0.401 |
|  | *p* | 0.467 | 0.531 |
| | $\eta^2_p$ | 0.018 | 0.014 |
|  | 90% CI | 0, 0.426 | 0, 0.406 |
| Condition<br>× | *F* | 0.339 | 0.319 |
|  | *p* | 0.565 | 0.577 |
| Context | $\eta^2_p$ | 0.012 | 0.011 |
|  | 90% CI | 0, 0.395 | 0, 0.391 |

<sup>#</sup>: The accuracy was subjected to the arcsine square root transformation before statistical testing.
<sup>##</sup>: 90% CI for Cohen's *f*.

**Table S3:** Two by two repeated ANOVAs of ERP components to the non-target Asian faces in Experiment 2.

| | | Context | Condition | Condition × Context |
| --- | --- | --- | --- | --- |
| N1 | $F$ | 1.525 | 0.619 | 0.094 |
| | $p$ | 0.227 | 0.438 | 0.762 |
| | $\eta^2_p$ | 0.050 | 0.021 | 0.003 |
|  | 90% CI <sup>##</sup> | 0, 0.519 | 0, 0.436 | 0, 0.323 |
| P2 | $F$ | 11.910 | 0.104 | 13.372 |
| | $p$ | 0.002 | 0.749 | 0.001<sup>#</sup> |
| | $\eta^2_p$ | 0.291 | 0.004 | 0.316 |
|  | 90% CI | 0.291, 0.939 | 0, 0.328 | 0.324, 0.979 |
| N2 | $F$ | 1.977 | 6.036 | 2.533 |
| | $p$ | 0.170 | 0.020 | 0.122 |
| | $\eta^2_p$ | 0.064 | 0.172 | 0.080 |
|  | 90% CI | 0, 0.551 | 0.125, 0.748 | 0, 0.585 |
| N170 | $F$ | < 0.001 | 1.004 | 0.059 |
| | $p$ | 0.993 | 0.325 | 0.810 |
| | $\eta^2_p$ | < 0.001 | 0.033 | 0.002 |
|  | 90% CI | 0, 0 | 0, 0.476 | 0, 0.297 |

<sup>#</sup>: Significant interaction effect. Simple effect analyses confirmed reliable RS effects on the P2 amplitudes to Asian faces in Species-Ctx ( $p=.043$ , RS effect=.698), but not in Race-Ctx ( $p=0.021$ , RS effect=-0.553). <sup>##</sup>: 90% CI for Cohen's $f$ .

**Table S4:** Behavior performance in the one-back task in Experiment 3.

|  |  | Accuracy (%) <sup>#</sup> | Reaction Time (ms) |
| --- | --- | --- | --- |
| Race-Ctx |  |  |  |
|  | Alt-Cond | 85±13.6 | 543±62.5 |
|  | Rep-Cond | 85±13.8 | 538±63.1 |
| Species-Ctx |  |  |  |
|  | Alt-Cond | 86±14.0 | 544±73.9 |
|  | Rep-Cond | 86±12.4 | 548±65.2 |
| ANOVA |  |  |  |
| Condition | *F* | 0.023 | 0.035 |
|  | *p* | 0.880 | 0.854 |
| | $\eta^2_p$ | 0.001 | 0.001 |
|  | 90% CI <sup>##</sup> | 0, 0.234 | 0, 0.261 |
| Context | *F* | 3.530 | 0.393 |
|  | *p* | 0.070 | 0.535 |
| | $\eta^2_p$ | 0.105 | 0.013 |
|  | 90% CI | 0, 0.629 | 0, 0.398 |
| Condition<br>×<br>Context | *F* | 1.964 | 0.554 |
|  | *p* | 0.171 | 0.463 |
| | $\eta^2_p$ | 0.061 | 0.018 |
|  | 90% CI | 0, 0.541 | 0, 0.421 |

<sup>#</sup>: The accuracy was subjected to the arcsine square root transformation before statistical testing.
<sup>##</sup>: 90% CI for Cohen's *f*.

**Table S5:** Categorization behavior performance in Experiment 3.

|  | Race-Context | Species-Context | Context Effect (paired t-test) |  |  |  |
| --- | --- | --- | --- | --- | --- | --- |
|  |  |  | *t*(29) | *p* | Cohen's *d* | 95% CI for mean difference |
| Accuracy <sup>#</sup><br>(%) | 93±5.7 | 94±7.3 | -1.268 | 0.215 | -0.23 | -0.11, 0.03 |
| Reaction<br>Time (ms) | 545±62 | 466±62 | 10.329 | <0.001 | 1.89 | 63.4, 94.6 |
| Performance<br>Efficiency<br>(RT/ACC) | 589±66 | 501±108 | 6.313 | <0.001 | 1.15 | 59.1, 115.8 |

<sup>#</sup>: The accuracy was subjected to the arcsine square root transformation before statistical testing.

